# Hierarchical population genetic structure and signatures of adaptation in *Lates calcarifer*

**DOI:** 10.1101/2024.06.30.600906

**Authors:** Matthew A. Campbell, Joy A. Becker

## Abstract

**Context:** *Lates calcarifer* is a widespread Indo-Pacific fish that is important in aquaculture, recreational and commercial fisheries. Genetic divergences from different data sources and sampling schemes have been reported.

**Aims:** To conduct phylogenetic and population genetic analyses from a geographically and phylogenetically representative data set to identify hierarchical divisions within *L. calcarifer*. We further test the evolutionary significance of genetic units in terms of signatures of adaptation.

**Methods:** Using a whole-genome sequence data set of 61 fish including an outgroup, we conducted phylogenetic and population genetic analyes. We also generated measures of F_st_, nucleotide diversity (π) and Tajima’s D (*D*).

**Key Results:** We identify three main lineages of *L. calcarifer* corresponding to the Indian subcontinent, Southeast Asia and Australasia. Subdivision within each of the three main lineages is also identified and characterized. Adaptively significant differences are indicated within and between the three main lineages.

**Conclusions:** *L. calcarifer* exhibits genetic divergences at different levels that originate before and during the Pleistocene. These divergences are associated with adaptive divergence but unclear phenotypic changes.

**Implications:** This study highlights the need for comprehensive sampling and integrative study of genotypes and phenotypes across the range of *L. calcarifer*.

## Introduction

Essential processes necessary for life on Earth are sustained by biodiversity, and biodiversity provides for human needs in various ways. Regardless of its limits and challenges, the basic unit for understanding biodiversity is the concept of species (e.g. Agapow et al., 2004). While the recognition of species is indeed important, so is the recognition of units below the species level that contribute to biodiversity. The ecological effects of intraspecific variation can meet our exceed species effect and therefore are important to document and recognize (Des Roches et al., 2018). Conservation of both species-level and intraspecific variation serves to promote ecological, cultural and economic functions of organisms, resilience and genetic variation for future adaptive potential (e.g. Layton et al., 2021; Lehnert et al., 2023; Schindler et al., 2010). Specific and intraspecific variation can be considered through both isolation and adaptation without the reliance on formal species taxonomy (Waples, 1991).

We apply these basic ideas of isolation and adaptation to identify variation to a Indo-Pacific *Lates*. Most Indo-Pacific *Lates* are classified under *Lates calcarifer* (Bloch 1790), and it is a valuable commercial taxon of cultural and recreational significance e.g. (Cluney, 2004; Edwards-Vandenhoek, 2015; Shamsi et al., 2020). The common names ‘barramundi’ and ‘Asian sea bass’ are widely used. A broad-ranging taxon, *L. calcarifer* is often considered to be distributed from the Persian Gulf eastwards to Taiwan and Australia, and into Oceania (**Figure 1**). Abed et al. (2023) note that there are no reliable reports from the Persian Gulf of native *Lates* and indicate a natural distribution through western India. Within this broader range, two regional endemics may be recognized, in Sri Lanka *L. lakdiva* Pethiyagoda & Gill 2012 and Myanmar *L. uwisara* Pethiyagoda & Gill 2012 (Pethiyagoda and Gill, 2012). We refer to all these fishes together as Indo-Pacific *Lates*.

**Figure 1.**
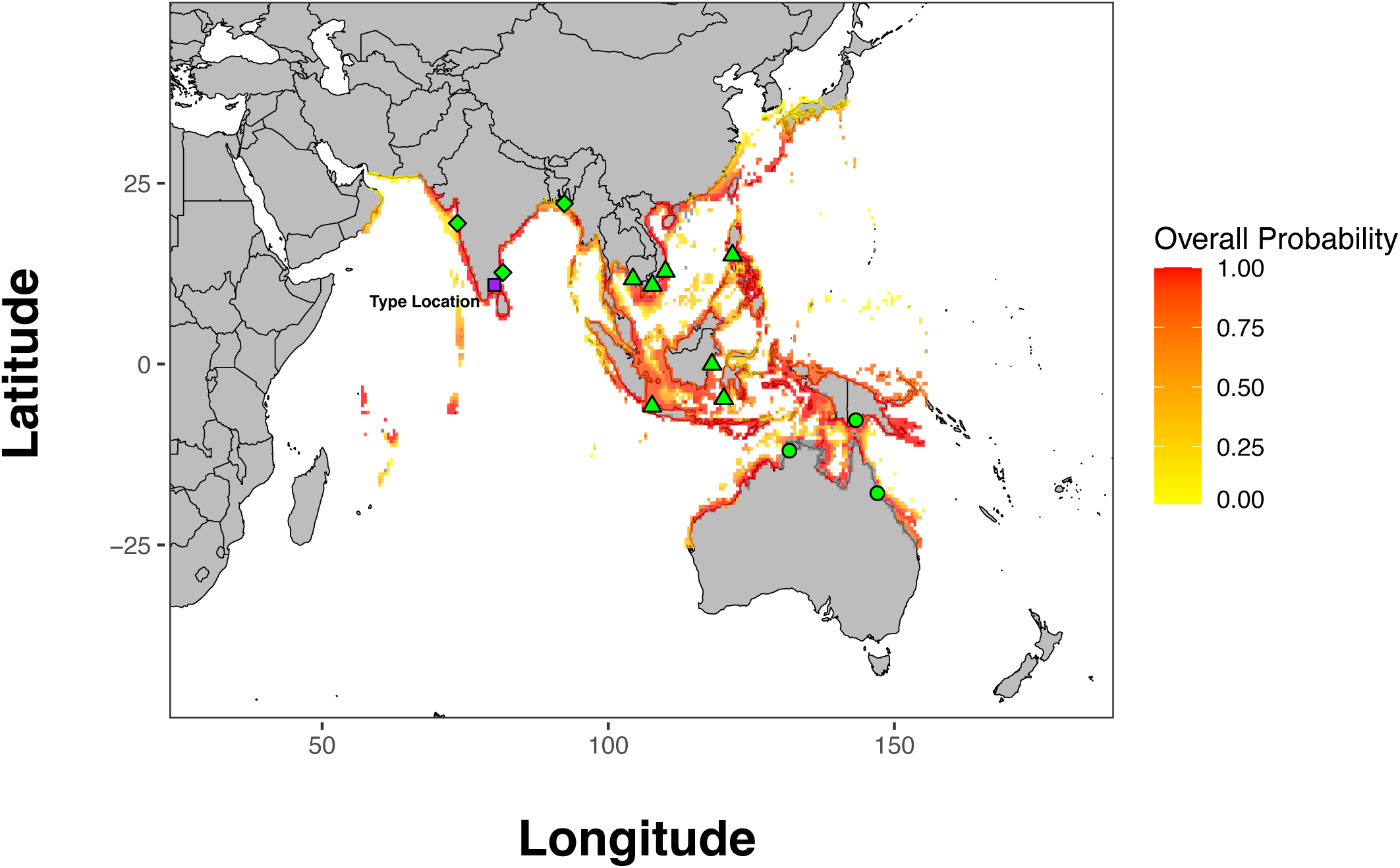
Distribution of *Lates calcarifer sensu lato* from AquaMaps (Kaschner et al., 2015) with probability of occurrence (Overall Probability) colored from yellow (low) to red (high). The type locality of *L. calcarifer* is indicated by a purple square. Sampling of main genetic lineages included in this paper are indicated by shape, AUS+NG circles, SEA triangles and IND diamonds. Locations of WGS are approximate as specific coordinates are not available. Cambodia is represented by four separate collection locations (shown as one).

Aquaculture, beginning in Thailand in the 1970’s, has increased greatly and Indo-Pacific *Lates* are now widely propagated across their range with some aquaculture outside the native range. Indo-Pacific *Lates* are now grown in diverse locations such as the United States of America (Arizona, Hawaii and Massachusetts), Israel, Taiwan and the Persian Gulf (Iraq) (Abed et al., 2023; Chen et al., 2024; Mathew, 2009). Propagation is also undertaken for stocking programs in lakes across northern Australia (e.g. Manton Dam, Northern Territory; Lake Kununurra, Western Australia) to support recreational fisheries and tourism in remote communities. Generally, Indo-Pacific *Lates* are considered to be protandrous hemaphrodites, catadromous, and tolerant of a wide range of salinities (Moore, 1979; Moore and Reynold, 1982). The sequential hermaphrodism of the taxon requires the importation of young males into aquaculture facilities as older fish become females, a potential source of hybridization and mixing of stocks of different origin. Movement for aquaculture, e.g. from Thailand to Taiwan also can result in the mixing of fish of different backgrounds and opportunities for release or escape (Chen et al., 2024). Other qualities of Indo-Pacific *Lates* such as high fecundity, ease of culture and rapid growth (i.e. harvestable size of 3 kg in less than two years) encourage their production (Rimmer, 2009).

The widespread culture and commercial fishing for Indo-Pacific *Lates* has led to genetic studies to understand population genetic structure. Initial mitochondrial barcoding revealed a deep divergence between Australian and Myanmar fish classified under *L. calcarifer* (K2P distance ∼ 9.50%) (Ward et al., 2008). Broader sampling geographically of mitochondrial lineages has substantiated this divergence, showing a distinctive clade of Indian subcontinent and Bay of Bengal fish (IND, including the Andaman Islands and Myanmar) separate from Southeast Asia (SEA, e.g. Singapore, Malaysia, Thailand and Indonesia) and Northern Australia (Vij et al., 2014). Vij et al. (2014) did include the sequences from Ward et al. (2008) and demonstrated that they are similar to a more broadly distributed IND mitochondrial lineage. Restriction site-Associated DNA sequencing (RADseq) applied to population-level samples of Indo-Pacific *Lates* (n = 131) from six localities (Thailand, Malaysia, Indonesia, Western Australia, Eastern Australia and Papua New Guinea) revealed a high degree of differentiation of SEA fish from Australia + New Guinea (AUS+NG). Pairwise F_st_ within SEA was 0.005-0.009 but 0.360-0.443 between SEA and AUS+NG without evidence of contemporary gene flow (Wang et al., 2016). Some investigation of population genomic structure was applied to a range-wide data set at a high level as part of a genome assembly study but did not investigate concordance between mtDNA and genomic DNA or population genetic metrics (Vij et al., 2016). Broadly, Vij et al. (2016) identified a separation of Indian Ocean fish, Southeast Asian fish and Australasian fish into genetic clusters. Population genetic structure range-wide examined with microsatellites (*K* = 4 genetic clusters presented) produced rather unclear distinctions of Indian Ocean fish from parts of Southeast Asia, and indicates a central Indonesian phylogroup and Australasian phylogroup (Jerry and Smith-Keune, 2014). No study to date, has analyzed both mitochondrial data and genomic data explicitly from the same individuals to evaluate concordance between data sources and resolve possible discordance between mitochondrial and genomic signals.

While there is evidence of genetic differentiation within Indo-Pacific *Lates* at a high phylogenetic level (Vij et al. 2014), this has only been evaluated thoroughly with molecular phylogenetic approaches using mtDNA. Within major genetic lineages, there is also genetic differentiation evident (e.g. Chenoweth et al., 1998; Shaklee and Salini, 1985; Yue et al., 2009), and the evolutionary significance of these differences are not substantiated. We apply a hierarchical approach to identify isolation (discreteness) of Indo-Pacific *Lates* genetic lineages with molecular phylogenetic and population genetic approaches, and test for adaptive significance between these lineages. Within lineages, we test for additional isolation and adaptive significance of these differences with population genetic approaches.(Lehnert et al., 2023). We further evaluate evidence of hybridization between main genetic lineages to determine if the hybridization is recent (F1, F2) and presumably a result of human activities or represents older admixture events.

## Methods

We approached the question of isolation within Indo-Pacific *Lates* by first constructing phylogenies using mitochondrial DNA and genome-wide SNPs. Subsequently, population genetic techniques were used, and divisions were examined at a high level (genetic lineages) and then within those lineages. We tested for evolutionarily significance through identifying possible adaptive loci between and within main lineages of Indo-Pacific *Lates.* An outstanding question is if hybridization is recent as a result of human mediated translocations or a result of natural admixture between lineages. We applied statistical testing to this question to determine if evidence of recent introgression in mtDNA was corroborated at the genetic level. Finally, we developed a timeline for diversification of Indo-Pacific *Lates* using a time-calibrated phylogenetic approach.

### Whole-Genome Sequence Data Set

Genome-wide sequence data from Indo-Pacific *Lates* were downloaded from the NCBI Sequence Read Archive under the BioProject accession PRJNA311498. The 61 samples in PRJNA311498 covered a broad distribution, from the Indian subcontinent, parts of Southeast Asia, Northern Australia and Papua New Guinea. Two additional sequences of BP were obtained from Bangladesh (PRJNA1021005) and for phylogenetic analyses an outgroup obtained (*L. japonicus,* n =1, PRJDB13763). The paired-end data was aligned to the BP reference genome GCF001640805.2 (TLL_Latcal_v3) with the Burrows-Wheeler aligner (BWA) with the mem algorithm (Li and Durbin, 2010, 2009). Alignments were sorted, properly paired reads kept and PCR duplicates removed with SAMTools (Danecek et al., 2021; Li et al., 2009). Total read counts, filtered read counts and depth of coverage were calculated from alignments with SAMTools. Those samples with coverage ≤ 3X were excluded from further analyses.

### Mitochondrial Phylogeny

A mitochondrial alignment from the WGS data was created by creating a Binary Alignment Map (BAM) file for each individual with more than 3X coverage only including reads aligned to the reference mitochondrial genome (NC_007439.1, samtools view). These BAM files were converted to consensus FASTQ files with the bam2cns subprogram of proovread (Hackl et al., 2014) then to FASTA files turning all bases with quality scores below 20 to N with seqtk (seq -a -q20 -n N) (https://github.com/lh3/seqtk). The mitochondrial Cytochrome Oxidase I (mtCOI) gene was extracted from the complete mitochondrial FASTA-formatted sequences with SAMtools (faidx NC_007439.1:6329-7879). For rooting, the corresponding region of the *L. japonicus* mitochondrial genome was extracted from the NCBI reference sequence (NC_034339.1) and sequences were aligned with MAFFT (Katoh et al., 2002; Katoh and Toh, 2008). A Maximum-Likelihood (ML) phylogeny of sequences was generated with IQTREE specifying a Hasegawa-Kishino-Yano model of nucleotide substitution with gamma-distributed rate variation (-m HKY+G) and support for nodes evaluated with rapid bootstrapping (-bb 10000) (Hoang et al., 2018; Nguyen et al., 2014). The tree was imported into R and visualized with the ggtree package (Yu et al., 2017) collapsing nodes to polytomies receiving less than 50% Bootstrap Support (BS) and rooting by the longest branch.

### Genome-Wide Phylogenetic and Network Analyses

We generated a genome-wide set of called SNPs with Analysis of Next Generation Sequencing Data (ANGSD) from the 24 main linkage groups in the BP assembly (Korneliussen et al., 2014). Quality of called SNPs was enforced by ensuring that a SNP was present in at least 90% of individuals, had a minimum Minor Allele Frequency (MAF) of 0.05, a minimum mapping quality of 20, minimum base quality of 20, a SNP *p* – val of 1.00 x 10^-6^ and a posterior cutoff of 0.90. A SAMTools model was specified and a PLINK-formatted file created as the output. We converted the PLINK-formatted file to a VCF-formatted file with PLINK (Purcell et al., 2007). Linked SNPS were removed with BCFTools +prune (-l 0.20 -w 10000).

The multispecies coalescent was applied to model gene-tree and species-tree discordance resulting from Incomplete Lineage Sorting, ILS (Maddison, 1997) on a set of called SNPs that included the outgroup *L. japonicus*. A species-tree was generated using SVDQuartets in PAUP* version 4.0a (Chifman and Kubatko, 2015, 2014; Swofford, 2003). Samples were pooled by sampling region for the species tree. A random set of quartets (evalQuartets=random), partially ambiguous SNPs were distributed over all compatible site patterns (ambigs=distribute), and 1,000 bootstrap replicates (bootstrap=standard nreps=1000) were options specified.

For ML phylogenetic analysis and network analysis we called SNPS with BP samples and a phylogeny of SNPs was inferred with IQTREE by identifying a model of nucleotide evolution with ModelFinder Plus and indicating the ascertainment bias correction should be applied (-m MFP+ASC) (Kalyaanamoorthy et al., 2017; Nguyen et al., 2011). IQTREE was used to further filter SNPs for suitability for ascertainment bias correction. Support for nodes was evaluated with 10,000 rapid boostraps (Hoang et al., 2018). As concatenated phylogenetic analyses do not explicitly model hybridization or ILS, we created an implicit network to identify potential hybridization or ILS in our data set with SplitsTree (Huson and Bryant, 2005). The same alignment provided to IQTREE was analyzed with SplitsTree.

### Population Genetic Structure and Statistics

Total signal in the data set was assessed by the generation of covariance matrix with ANGSD and a PC analysis. The covariance matrix was generated by with a single read sampling approach due to a wide range of individual coverage (minimum 4.73, maximum 34.81, mean 9.38 and median 6.41) with the options -doIBs 1 and -doCov 1 in ANGSD. The 24 main linkage groups of the reference assembly were sampled for this analysis. A variable site was required to be present in >90% of individuals with a minimum MAF of 0.05. A minimum mapping quality of 10 was specified with a minimum base quality of 20 and a GATK genotype likelihood model was used (-GL 2). The resulting covariance matrix was imported into R for PC analysis.

To calculate genetic differentiation in terms of F_st_ was calculated by generating Site Frequency Spectra (SFS) for each sampling region with ANGSD and the realSFS subprogram, splitting Indonesia into two locations based on metadata available with the sequences (“K” n=5 and and “SJ” n=5). Vietnam (n=2), Indonesia “SU” (n=1) and Bangladesh (n=2) were excluded due to small sample sizes. Two-dimensional SFS were calculated between each possible pair of sampling regions and F_st_ calculated (see code repository accompanying this paper). Nucleotide diversity (*π*) and Tajima’s D (*D*) estimates were calculated for each chromosome with realSFS using the saf2theta option followed by the thetaStat subprogram of ANGSD and the do_stat option.

Admixture analysis was conducted with NGSadmix (Skotte et al., 2013) on genotype likelihoods generated with ANGSD as for the PC analysis using a single read sampling approach. The resulting BEAGLE-formatted file was used with NGSadmix to produce an admixture plot of *K* = 2-5 genetic clusters.

### Tests for Adaptative Genetic Variation

Genotype calls produced for phylogenetic network analysis after pruning were imported into R for analysis with PCAdapt (Luu et al., 2017; Privé et al., 2020). The *pcadapt* function was then used to generate a scree plot and identify an objective number of PCs with the first 20 PCs examined. From the resulting object from the *pcadapt* function, outlier loci were identified by filtering for a Benjamini and Hochberg (1995) and Bonferroni corrected *p* – values with significance values less than 0.10. We repeated this analysis for individuals in each main genetic lineage by filtering for variants with a minimum minor allele frequency of 0.05 and examining the first 5 PCs.

### Evaluation of Hybridization Levels

It is possible that Southeast Asian *Lates* exhibit a mixture of hybridization levels between IND and SEA lineages, with possible F1s, F2s and advanced backcrosses. Two wild-caught samples in the data set from Cambodia (sequence read accessions SRR3165998, SRR3165999) had IND lineage mtDNA sequences (see results, Figure 2). We applied NewHybrids (Anderson and Thompson, 2002) to test if these two Cambodian fish were recent (F1, F2) or older hybrids (advanced backcrosses). Using the called and pruned SNP data sets, we excluded recently reported potential chromosomal inversions (Campbell and Hale, 2024) and imported the data into R and analyzed with functions of the dartR package (Gruber et al., 2018). We restricted the analysis to fish from the SEA and IND lineages as NewHybrids functions with two populations and mixtures of those populations then filtered out low MAF SNPS (threshold of 0.25). The *gl.nhybrids* function of dartR was used to drive a NewHybrids analysis with India Western Coast and Bangladesh sampling locations designated as one parental lineage (P0) and Indonesia-K and Indonesia-SU as another (P1). We allowed SNPs to not be entirely fixed between parental populations (threshold=0.01) and chose a subset of 200 SNPs for analysis with the ‘AvgPIC’ criterion. Analysis included a 100,000 step burnin and 50,000 sweeps.

**Figure 2.**
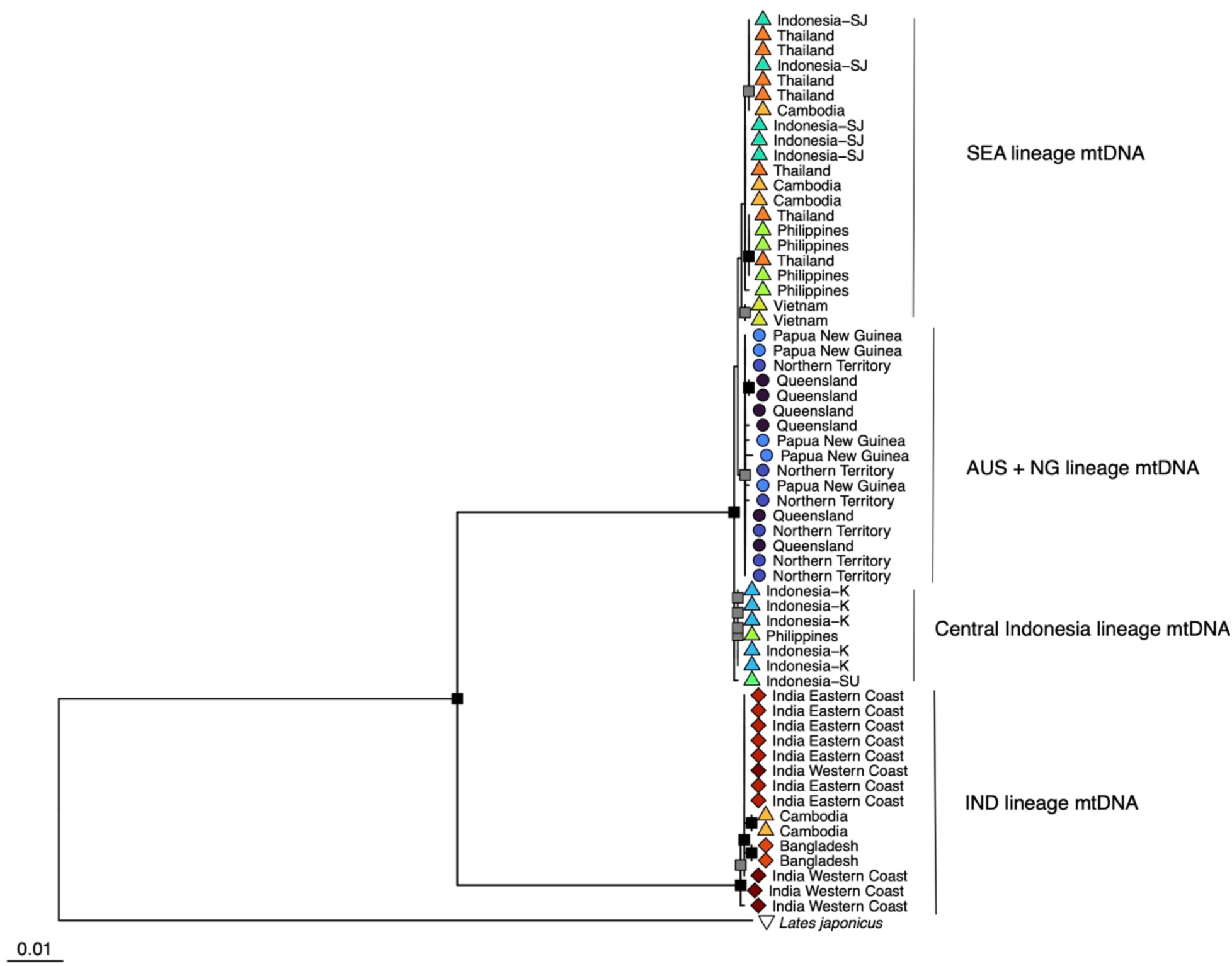
Maximum-Likelihood phylogeny of mitochondrial Cytochrome Oxidase I from 60 Indo-Pacific *Lates* individuals derived from whole-genome sequence data. The tree is rooted by *L. japonicus*. Node shape indicates the putative lineage of origin and collection information is provided as tip labels. Nodes with < 50% bootstrap support are collapsed to polytomies. Nodes with ≥ 75% bootstrap support but < 90% bootstrap support are indicated as grey squares and nodes ≥ 90% bootstrap support as black squares.

### Divergence Time Estimation

Existing efforts have covered a broad geographic and lineage sampling of Indo-Pacific *Lates* with barcoding (mitochondrial *cox1*) (Vij et al., 2014; Ward et al., 2008). The effort of this study was then to calculate divergence times of the three lineages (IND, SEA and AUS+NG). We downloaded barcode sequences from the Barcode of Life Database and Genbank for Latidae. With BLAST (blastn) we confirmed the major lineages of Indo-Pacific *Lates* in barcode data (Altschul et al., 1997) then selected representative sequences from these three lineages to conduct a divergence time analysis (**Table 1**). We also searched for representation across Latidae for broad taxonomic representation of this family including sequences from *L. japonicus*, *L. microlepis* and *L. niloticus*. We included representatives of the two main mitochondrial lineages present in *Psammoperca* corresponding to *P. waigiensis* and *P. datnioides*. Centropomidae has been confirmed to be the sister lineage of Latidae in molecular phylogenetic study e.g. (Campbell et al., 2013; Chen et al., 2007; Li et al., 2011) and inclusion of Centropomidae allows dating the Time to Most Recent Common Ancestor (TMRCA) of Centropomidae and Latidae to 48.6 million years ago (mya) (Campbell et al. 2013; Otero, 2004). We also included sequences from the outgroup lineages Channidae and Nandidae as an additional calibration point dating the TMRCA of those taxa to be 48 mya (Santini et al., 2009).

**Table 1.** Sample origins by region of whole-genome sequence data set for Indo-Pacific *Lates*. The sample size and average coverage is presented for each sampling region.

| Region | Additional Geographic Information | Source | Sample Size | Average Coverage |
| --- | --- | --- | --- | --- |
| Northern Territory, Australia | Darwin Harbor | Wild-Caught | 6 | 5.80 |
| Queensland, Australia | Hinchinbrook Channel | Wild-Caught | 6 | 5.34 |
| Papua New Guinea | Fly River | Wild-Caught | 5 | 5.83 |
| Indonesia | K - Kalimantan | Unknown | 5 | 5.73 |
| Indonesia | SJ – South of Jakarta | Wild-Caught | 5 | 17.8 |
| Indonesia | SU - Sulawesi | Unknown | 1 | 6.13 |
| Philippines |  | Hatchery Broodstock | 5 | 6.32 |
| Vietnam |  | Hatchery Broodstock | 2 | 6.21 |
| Cambodia | Three locations | Wild-Caught | 5 | 5.90 |
| Thailand | East Coast | Wild-Caught | 7 | 15.60 |
| Bangladesh | Bakkhali River | Unknown | 2 | 10.04 |
| India Eastern Coast | Tamil Nadu | Wild-Caught | 7 | 8.50 |
| India Western Coast | Mumbai | Wild-Caught | 4 | 20.22 |

**Table 2.** Taxa and sequences included in time-calibrated phylogeny of Indo-Pacific *Lates* mitochondrial lineages. Lates *calcarifer* is represented by three lineages, Australia + New Guinea (AUS+NG), Southeast Asia (SEA) and Indian subcontinent (IND) as described in the text. The family is given, with the taxon, corresponding BOLD or NCBI accession and the geographic origin of the sample if known.

| Family | Taxon | Accession | Origin |
| --- | --- | --- | --- |
| Latidae | *Lates calcarifer* AUS+NG | EU189379.1 | Western Australia |
|  | *Lates calcarifer* SEA | KJ573921.1 | Indonesia |
|  | *Lates calcarifer* IND | KJ573895.1 | Andaman and Nicobar Islands |
|  | *Lates japonicus* | LC269832.1 | Japan |
|  | *Lates microlepis* | ISZA082-21 | Democratic Republic of the Congo |
|  | *Lates niloticus* | LT853908.1 | Not Specified |
|  | *Hypopterus macropterus* | LC269834.1 | Australia |
|  | *Psammoperca waigiensis* | OQ387793.1 | Philippines, Luzon |
|  | *Psammoperca datnioides* | NC035737.1 | Australia, Darwin |
| Centropomidae | *Centropomus ensiferus* | MW183492.1 | Brazil, Para, Braganca |
|  | *Centropomus undecimalis* | JQ841102.1 | Belize, Belize City |
| Channidae | *Channa maculata* | KP112195.1 | China, Hubei, Wuhan |
| Nandidae | *Nanus nandus* | KX245097.1 | India |

Sequences were aligned with MAFFT (Katoh et al., 2002; Katoh and Toh, 2008), in frame coding was confirmed by translation in the NCBI ORF finder and a NEXUS file made for a time-calibrated phylogenetic analysis in MrBayes (Ronquist et al., 2012). The third codon position was excluded due to the age of the data set and expected saturation occurring in mitochondrial sequences from carangimorph taxa at these timescales e.g. (Campbell et al., 2014). A nucleotide substitution model of General Time Reversible (GTR) with a proportion of invariant sites (I) and gamma-distributed rate variation (Ι′) was specified. Constraints were applied to Centropomidae + Latidae to date with an exponential distribution used, offsetexp(48.60, 63.18), and the outgroup, offsetxp(48.00,62.40). Initially, three runs and three chains were specified with a run length of 1 million generations sampled ever 1,000 generations and a 25% burnin applied. Evaluation of parameters and convergence with Tracer (Rambaut et al., 2018) was assessed, and run length increased until sufficient ESS for all parameters (>200) was reached. Tree summarization was done in MrBayes with the sumt command and an all compatible consensus tree made.

## Results

### Whole-Genome Sequence Data Set

From the original 63 Indo-Pacific *Lates* sequences, 60 had coverage over 3X and were included in additional analyses (**Table 1**). Sample sizes for regions were 13 for putative members of the IND lineage, 30 for putative members of the SEA lineage, and 17 for putative members of the AUS+NG lineage from ten sampling regions (**Figure 1**). The *L. japonicus* sequence aligned to the *L. calcarifer* reference genome with high success with 110X coverage and was down-sampled to 20X coverage with SAMtools to be more consistent with genome coverage measures of Indo-Pacific *Lates*.

### Mitochondrial Phylogeny

The 61 samples (60 Indo-Pacific *Lates* and one *L. japonicus*) produced an alignment of 1,576 base pairs in length with 71 distinct patterns and 128 parsimony-informative sites. In the alignment, 1,441 sites were invariant. The ML phylogeny indicates a clear division of *L. calcarifer* IND individuals with two Cambodian-origin fish from all other fish (**Figure 2**). Support for monophyly of this IND mtDNA lineage is maximal. Support for monophyly of SEA + AUS+NG is maximal, and support for a clade of *L. calcarifer* AUS+NG is moderate (BS=80%). Individuals sourced from the putative *L. calcarifer* SEA clade do not form clear geographic clusters, with the exception of a Central Indonesia mtDNA lineage that receives low support for monophyly (BS=57%) and includes a single sequence from the Philippines. Support for the arrangement of major mtDNA lineages SEA and AUS+NG lineages is low (BS=55%). The Assemble Species by Automatic Partition algorithm indicated the strongest support for two species, with a threshold distance of 4.10% (results provided in the Data Supplement). This division separates the IND mitochondrial lineage from the combined SEA and AUS+NG lineage.

### Genome-Wide Phylogenetic and Network Analyses

For analysis with SVDQuartets to generate a rooted species-tree, 8,529 variants that all have unique alignment patterns were analyzed and is presented as **Figure 3**. The SNP alignment consisted of 6,456 parsimony informative and 2,073 singleton sites. Three lineages were apparent in the species-tree with maximal support for monophyly corresponding to the *L. calcarifer* IND, *L. calcarifer* SEA and *L. calcarifer* AUS+NG. The SEA and AUS+NG lineages were highly supported (BS=100%) as sister lineages. Within the IND lineage, the India Eastern Coast and Bangladesh sampling locations were indicated to be sister lineages (BS=100%). The AUS+NG lineage individuals from the Northern Territory and Papua New Guinea were indicated to be most closely related (BS=100%). The first division within the SEA lineage is between Central Indonesia sampling locations (Indonesia-SU + Indonesia-K) and receives maximal support for monophyly and branching patterns. The Philippines sampling location separates with the next successive branching within the SEA lineage (BS=100%). Relationships among Cambodia, Indonesia-SJ, Thailand and Vietnam sampling locations were not clearly resolved.

**Figure 3.**
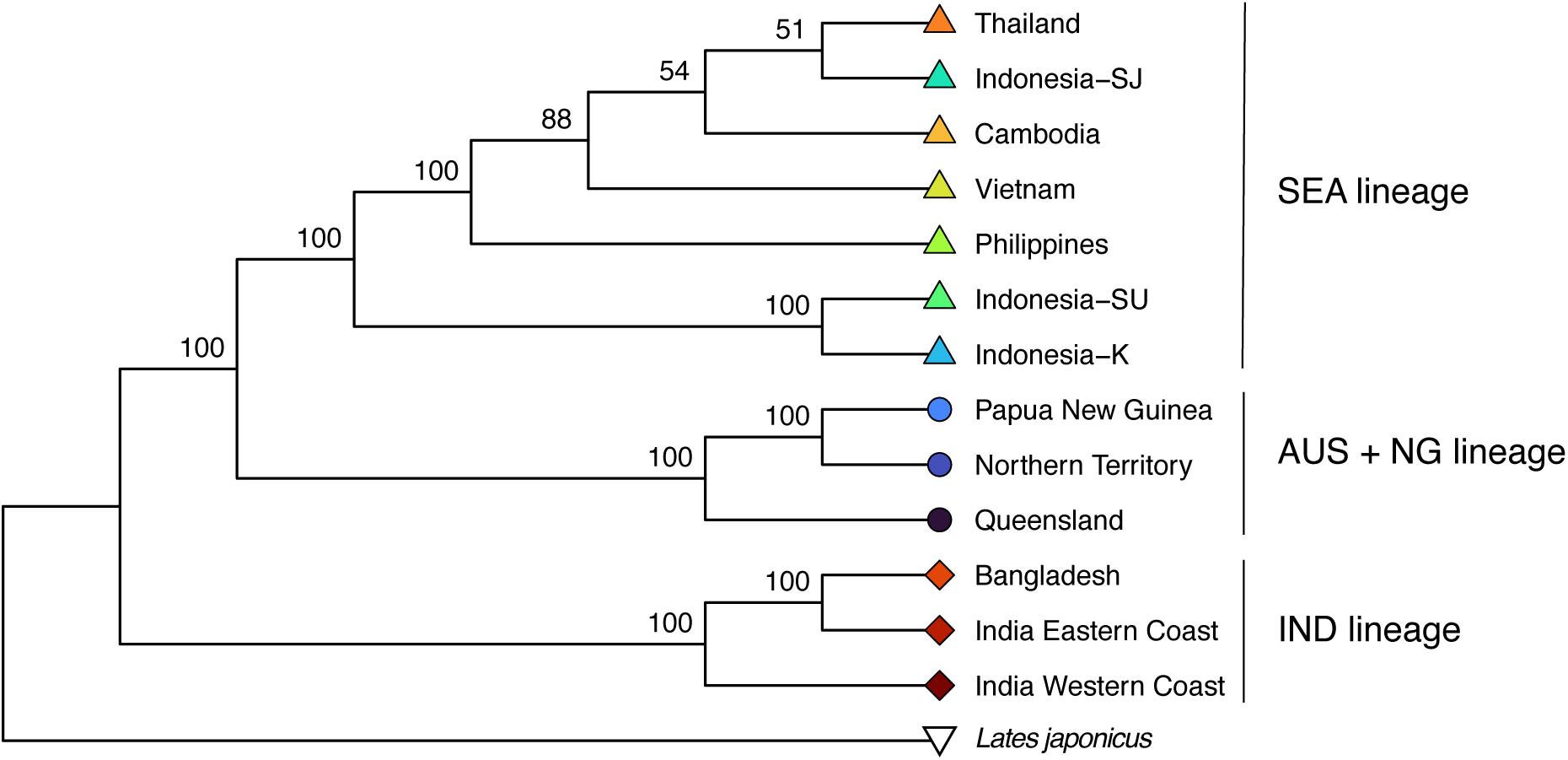
Species-tree from SVDQuartets of Indo-Pacific *Lates* sampling locations in this study based on 8,529 variants rooted by *Lates japonicus*. Values at nodes are bootstrap support values.

For ML phylogenetic analysis and phylogenetic network analysis an alignment of 60 BP individuals with 7,661 SNPs having 7,661 distinct unique alignment patterns and 5,571 parsimony informative sites was created. The alignment contains 2,090 singleton sites. Support for monophyly of IND lineage fish is maximal, as well as geographic divisions into Eastern and Western Indian subcontinent (**Figure 4A**). Indo-Pacific *Lates* from SEA as a grouping receives low support (BS = 76%) with concordance between sampling locations and placement not complete. Indonesia-K and the Philippines sampling locations were highly supported to be monophyletic, but Cambodia, Thailand and Indonesia-SJ were not. Support for a group of AUS+NG sampling locations is maximal, with maximal support for division between Queensland and Northern Territory + Papua New Guinea fish. Monophyly of samples from the Northern Territory is poorly supported (BS = 68%).

**Figure 4.**
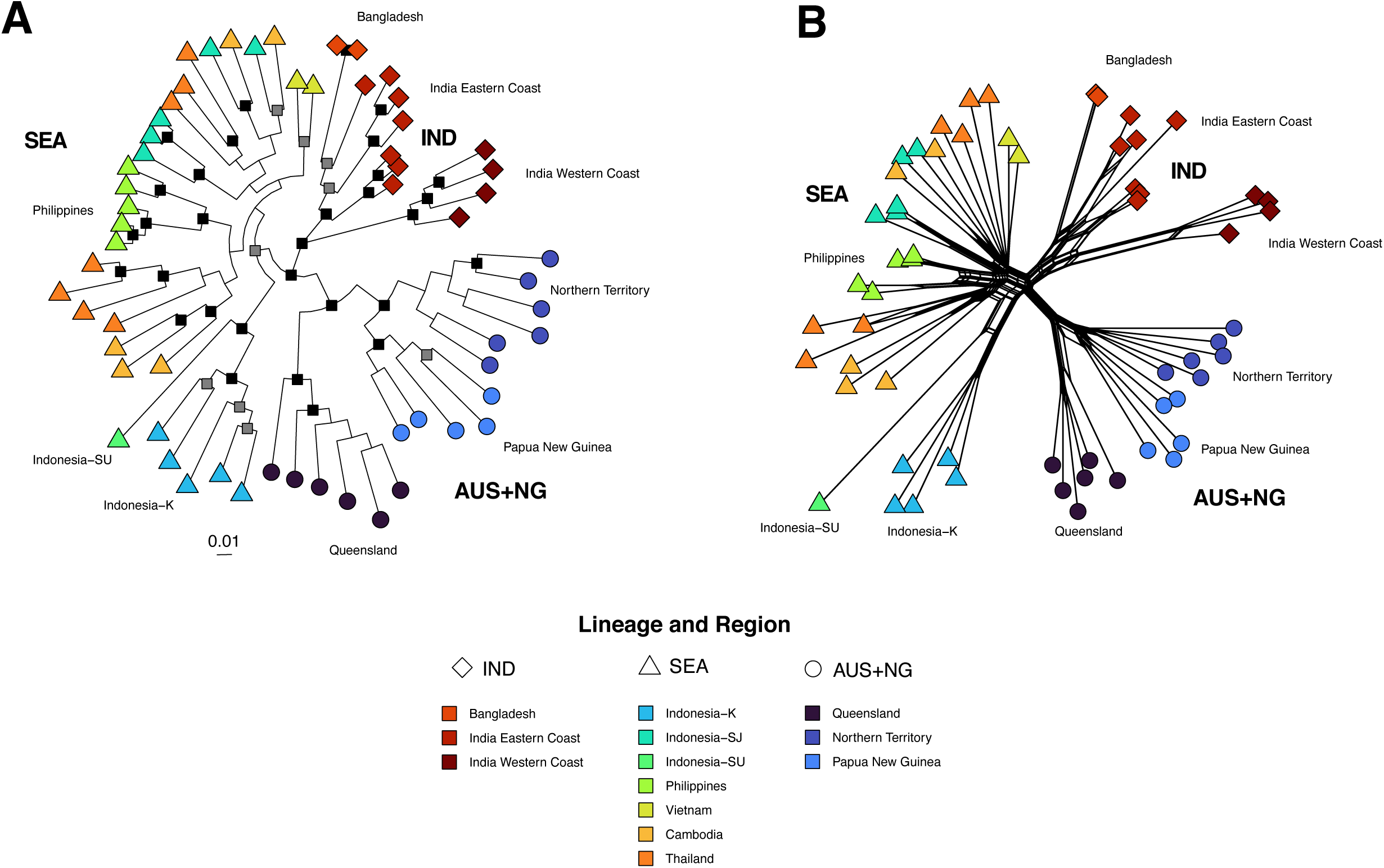
(A) Maximum-Likelihood phylogeny of genome-wide SNPs of 60 individual Indo-Pacific *Lates*. Nodes with < 75% bootstrap support are collapsed to polytomies. Nodes with ≥ 75% bootstrap support but < 90% bootstrap support are indicated as grey squares and nodes with ≥ 90% bootstrap support as black rectangles. **(B)** Implicit phylogenetic network of genome-wide SNPs from 60 individual Indo-Pacific *Lates*. Node shape indicates the putative lineage of origin and color indicates collection location.

The phylogenetic network resolves an IND lineage containing two sub-lineages (India Eastern Coast + Bangladesh, India Western Coast) with separation of all three sampling locations (**Figure 4B**). The SEA lineage is present separated between Central Indonesia and all other locations as two main sub lineages, with reticulations with the IND lineage occurring with a subgroup of 10 individuals, disrupting monophyly of some sampling locations. The Indonesia-K and Philippines locations were monophyletic, but the Cambodia, Thailand and Indonesia-SJ sampling locations were not. The AUS+NG grouping is composed of three subclades Queensland (Australia), Northern Territory (Australia) and Papua New Guinea. In the network, Northern Territory and Papua New Guinea fish were most-closely related to each other.

### Population Genetic Structure and Statistics

The PC analysis included 7,321,852 variable sites in the genome and indicated a separation into three larger genetic clusters corresponding to IND, SEA and AUS+NG individuals (**Supplemental Figure S1**). The first PC separated AUS+NG, SEA and IND individuals (PC 1 12.57% of variation) with the second PC showing variation within SEA and IND clusters (PC2 4.71%). Clear separation of Indian East Coast + Bangladesh and Indian West Coast individuals is present on both PC 2 and PC 3 (PC 3 3.51%).

Genetic differentiation (F_st_) was lowest within main genetic lineages, such as Thailand and Cambodia (F_st_ = 0.06) within the *L. calcarifer* SEA lineage and between the Indian East Coast and Indian West Coast sampling locations (F_st_ = 0.11) within the *L. calcarifer* IND lineage (**Figure 5A**). The highest levels of genetic differentiation were found between main genetic lineages, between AUS+NG and SEA F_st_ = 0.15-0.35, between AUS+NG and IND F_st_ = 0.22-0.33, between SEA and IND F_st_ = 0.12-0.26). Nucleotide diversity is the lowest in the AUS+NG lineage (*π* = 20,067-22,775) and highest in the IND lineage (*π* = 207,299-214,485) (**Figure 5B**). Tajima’s D was strongly negative (< -1) in the Indonesia-SJ, Thailand, India Eastern Coast and India Western Coast sampling locations (**Figure 5C**). The AUS+NG sampling locations had *D* values between -0.58 and -0.77.

**Figure 5.**
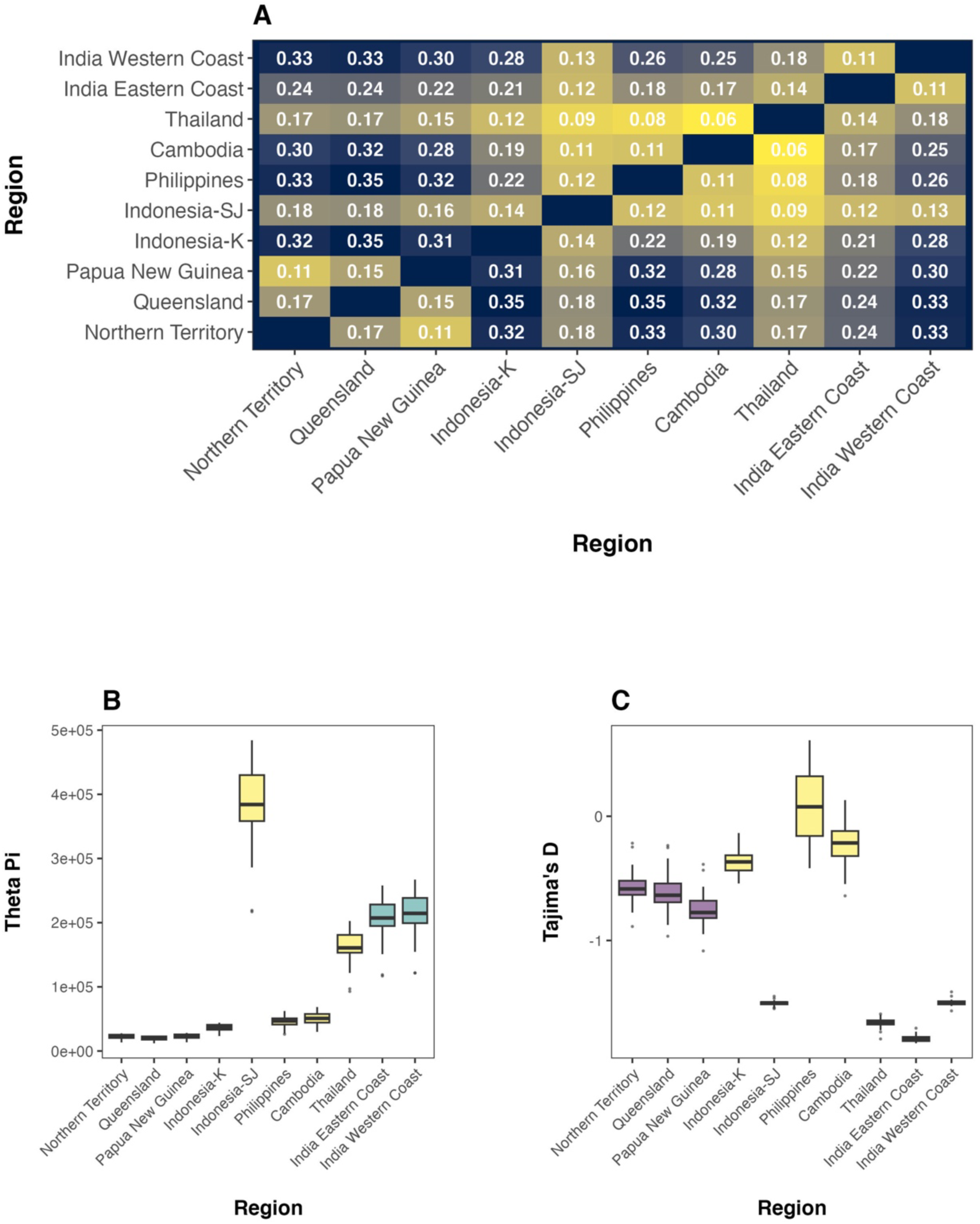
Population genetic statistics of Indo-Pacific *Lates* from WGS data. **(A)** Pairwise Fst values between regions. **(B)** Nucleotide diversity of sampling locations in the WGS data set. **(C)** Tajima’s D of sampling locations in the WGS data set. Indonesia is represented by two sampling locations. Indonesia-SU, Vietnam and Bangladesh were excluded due to low sample sizes (n ≤ 2).

Admixture analysis of 7,321,852 genotype likelihoods from 60 BP individuals for *K* = 2-5 is presented as **Supplemental Figure S2**. At *K =* 2, the Bangladesh, Indian Eastern Coast and Indian Western Coast have high ancestry coefficients (*Q*) > 0.99 for a single genetic cluster and the AUS+NG lineage and Indonesia-K sampling location have high ancestry coefficients for the other. The remaining SEA sampling locations have a mixture of contributions from both genetic clusters. At *K* = 3 the AUS+NG individuals form a clearly genetic cluster with *Q* > 0.99 for ancestry coefficients, the SEA sampling locations were heterogenous mixture proportions from sampling locations largely composed of the third genetic cluster, with the Indonesia-K and Indonesia-SU sampling locations exhibiting a near uniform ∼0.27% admixture proportion with the genetic cluster typical of AUS+NG sampling locations. Higher values of *K* lead to separating of genetic units within the SEA lineage that may or may not correspond to sampling locations.

### Tests for Adaptation

The variation within the pruned genotype calls of 22,088 SNPs from the n=60 WGS sequences with >3X coverage supported an optimal *K* = 3 PCs (**Supplemental Figure S3**). These PCs separated the three main lineages of *L. calcarifer*, with the third PC separating the Indonesia-K sampling location and to a lesser degree the Indonesia-SU individual from other SEA fish. Benjamini and Hochberg (1995) adjusted *p* – values (BH) identified 134 candidate adaptive SNPs and Bonferroni corrected *p* – values (BF) indicates 34 candidate adaptive SNPs across the three PC axes.

Filtering to IND individuals (n=13) resulted in 12,075 SNPs with a minimum MAF of 0.05, with the first PC separating the Bangladesh, India Eastern Coast and India Western Coast with 738 BH and 379 BF candidate adaptive SNPs identified on this axis (**Supplemental Figure S4**). The second PC axis separated India Eastern Coast with 285 BH and 59 BF candidate adaptive SNPS. From the SEA lineage, 30 individuals with 17,556 SNPs with a minimum MAF of 0.05 were analyzed. The first three PCs were considered, separating Indonesia-K and Indonesia-SU sampling locations on PC 1, the Philippines along PC 2 and some Thailand and Cambodia individuals along PC 3 with 190 BH and 29 BF candidate adaptive SNPs identified in total (**Supplemental Figure S5**). The data set of AUS+NG individuals (n=17) for loci with a minimum MAF of 0.05 resulted in 14,642 SNPs. We found strong support for *K* = 2 PCs, the first PC separating Queensland from the Northern Territory and Papua New Guinea (**Supplemental Figure S6**). The second PC separated Papua New Guinea from the other locations with 234 BH candidate adaptive SNPs and 57 BF candidate adaptive SNPs across the two PC axes.

### Evaluation of Hybridization Levels

The NewHybrid analysis of 43 individuals from the SEA and IND lineages indicated all IND lineage fish were most likely a parent population (P0, **Supplemental Figure S7**). This includes India Eastern Coast, which was not designated *a priori* as a potential parent population. Indonesia-K and Indonesia-SU were resolved as P1 fish with high probability. The two wild-caught fish from Cambodia with IND lineage mtDNA were indicated to be F2s, with other fish from Cambodia P1 (n=2) and a backcrossed P1 fish.

### Divergence Time Estimation

We compiled a barcode alignment of 13 sequences with 660 characters resulting in 440 characters (399 invariant) when including the first and second codon positions only. Sufficient effective sample sizes (ESS >200, range of 712-3,002) was reached with 20 million generations and two runs of three chains each as evaluated after a 25% burnin. The resulting consensus tree presented in **Figure 6**. The median divergence time of the IND lineage from the other two lineages is estimated to be 6.76 mya with a 95% Highest Posterior Density (HPD) of 1.31-14.24 mya. The divergence of AUS+NG from SEA is estimated to be 1.02 mya with a 95% HPD range of 0.00-5.03 mya. The relationships of Indo-Pacific *Lates* mitochondrial lineages were highly supported, with the placement having a Posterior Probability (PP) of 0.99, and the sister relationship between AUS+NG and SEA having a PP of 0.97. The NEXUS file for use with MrBayes and the consensus tree are provided in Data Supplement of this paper.

**Figure 6.**
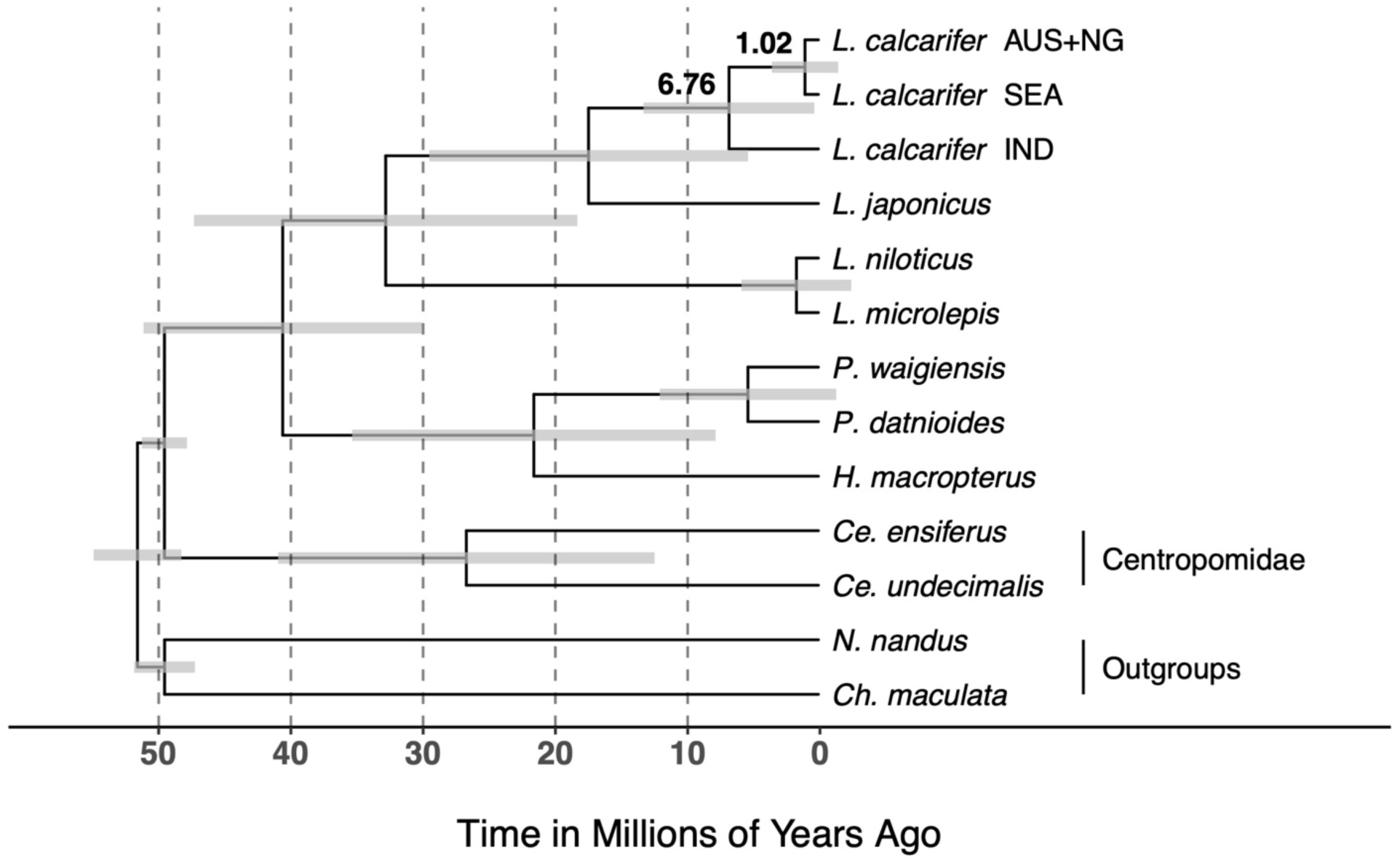
Time-calibrated phylogeny of main mitochondrial lineages of Indo-Pacific *Lates*. Node age is from the posterior medians with 95% Highest Posterior Density (HPD) indicated by horizontal bars.

## Discussion

We identify from mitochondrial and nuclear genomic data three main genetic lineages of Indo-Pacific *Lates*. These are an Indian Ocean lineage (IND), and a Pacific Ocean lineage split between Southeast Asia (SEA) and Australasia (AUS+NG). A Central Indonesian genetic cluster, representing Borneo and Sulawesi appears as distinctive group with placement poorly supported in our mtDNA analysis (Figure 2). Phylogenetic and population genetic analyses using data from the nuclear genome, place the Central Indonesian samples as more closely related to Southeast Asian fish (SEA), and we treat them as part of that (SEA) lineage.

The three main genetic lineages of Indo-Pacific *Lates* are present with substantial differences quantified in terms of genome-wide estimates of F_st_, candidate adaptive loci and deep divergence times. We observed F_st_ values up to 0.36 and in terms of divergence times, we estimated 6.76 million years for the divergence of IND from other Indo-Pacific *Lates* and a divergence time estimate of 1.02 million years for AUS+NG from SEA. The previously published estimate based on RADseq data for divergence times by Wang et al. (2016) is restricted to AUS+NG from the SEA lineage and is an approximate 644,100-878,900 years before present. Both of these estimates, the estimate presented here from fossil-calibrated phylogeny and Wang et al. (2016), support a divergence during the glacial cycling and changes in sea level and conditions during the Quaternary. The three main lineages of Indo-Pacific *Lates* exhibit discreteness (genetic isolation) and evolutionary significance (i.e. adaptive differentiation).

Within each of the three main lineages, there was additional evidence for discreteness and evolutionary significance. The IND sampling locations were divided phylogenetically (**Figures 3 & 4**), and in PC analysis (**Supplemental Figure S4**) with candidate adaptive SNPs identified. We find that all three sampling locations within IND (Bangladesh, India Eastern Coast and India Western Coast) can be separated clearly with genomic data but exhibit some mixing of mtDNA haplotypes. The SEA lineage exhibited notable separation of the Indonesia-K and Indonesia-SU sampling locations, the Philippines sampling location and some Thailand fish (**Figure 4**, **Supplemental Figure S5**). Clear differentiation of all Philippines and Thailand fish may be impacted by hybridization (**Supplemental Figure S2**). The RADseq data set of Wang et al. (2016) did not show clear evidence of genetic structure with three sampling locations within SEA, but broader sampling with microsatellites did appear to support genetic structuring (Yue et al., 2009). The AUS+NG sampling exhibited a clear division between Queensland and other sampling locations (**Figures 3 & 4**) and all three sampling locations could be clearly differentiated on two PC axes with candidate adaptive loci apparent (**Supplemental Figure S6**). Within the AUS+NG lineage there are several studies demonstrating genetic differentiation within that lineage (e.g. Chenoweth et al., 1998; Loughnan et al., 2019; Shaklee and Salini, 1985; Wang et al., 2016) with 16 discrete genetic populations identified in Australia only (Keenan, 1994).

### What is Lates calcarifer?

Our fundamental understanding of nature is based at the species level and the lack of recognition of species is a major knowledge shortfall in conservation and management (Brown and Lomolino, 1998). Molecular genetic techniques have continued to identify candidate species of fishes which may or may not have diagnostic morphological characters typical of species descriptions (e.g. Campbell et al., 2023; Moyle and Campbell, 2022). Instead of being a historical aspect of biology, the need for continued taxonomic work is very important (e.g. Delić et al., 2017; Godfray et al., 2004). Taxonomic work often involves the detailed anatomical investigation of organisms that can reveal phenotypic differences related not only to speciation, but also life history variation. What remains outstanding is if Indo-Pacific *Lates* can be accurately represented within Linnean taxonomy or if an alternative approach is better suited to the biological realities of Indo-Pacific *Lates*.

The most recent taxonomic effort relevant to BP led to the description of *L. lakdiva* Pethiyagoda & Gill 2012 from western Sri Lanka and *L. usiwara* Pethiyagoda & Gill 2012 from eastern Myanmar (Pethiyagoda and Gill, 2012). The specimens of *L. usiwara* were the same as featured in the barcoding study of Ward et al. (2008) highlighting deep mitochondrial divergences, but in a wider context the barcode sequences of *L. usiwara* are identical to samples from the Indian subcontinent (Vij et al., 2014). However, in comparison to *L. calcarifer*, a differential diagnosis was possible and the distribution of *L. usiwara* is perhaps indicative of a recently derived species or ecotype. The anatomical study undertaken by Vij et al. (2014) along with barcoding did not clearly separate all these newly described species of *Lates* from *L. calcarifer* noting small sample sizes used in the description of *L. usiwara* and *L. lakdiva* and conclude that there is substantial mismatch between anatomical descriptions of new species of *Lates* (within the Indian Ocean) and candidate species of *Lates* apparent genetically (between the Indian and Pacific oceans at the broadest scale). Barcode sequences, it should be noted, will not reflect proportion of ancestry as do genomic data and ascertaining the ancestry proportions of fish used in taxonomic studies is critical going forward due human translocations of *Lates*. Furthermore, it is possible for a high degree of genome-wide differentiation or substantial reproductive isolation be present that would not be apparent in barcode data. Pethiyagoda and Gill (2012) themselves observe that due to the large body size and sequential hermaphodism of *L. calcarifer sensu lato*, they were restricted to examining many juveniles for their study.

The three genetic lineages of Indo-Pacific *Lates* remain unclear in distribution and have unclear anatomical differentiation (Vij et al., 2014). Portions of the range, such as Taiwan, remain unsampled for genomic data and genetic structuring is apparent across all three lineages (**Figures 3 & 4, Supplemental Figures S1-S6**). Differentiation between and within the main lineages can be clarified with additional sampling and representation of nominal taxa in genomic data sets (i.e. *L. usiwara* and *L. lakdiva*). Given the strong genetic differentiation and deep inferred time scales (**Figure 6**), one possible resolution is to recognize at least two species-level taxa that correspond to the IND lineage and a combined SEA and AUS+NG lineage. At a potential species-level, the taxonomy of Indo-Pacific *Lates* is complex and the recognition of types and their origins is also challenging, reviewed by Pethiyagoda and Gill (2012). The type locality of *L. calcarifer* (Bloch 1790) may be Tamil Nadu, India (10.4°N, 79.3°E) (Pethiyagoda and Gill, 2012). Thus, the IND lineage may be represented by this name. There are numerous available names that may be applicable to the SEA lineage, but this requires substantial time and expertise to resolve. There is, however, an available name, *L. cavifrons* (Alleyne & Macleay 1877) that could be applied to AUS+NG fish. The collection locality is either the Torres Strait or the coast of New Guinea with a lost holotype with the drawing providing the current synonymy (Pethiyagoda and Gill, 2012).

Without formal taxonomic recognition, the three main genetic lineages of Indo-Pacific *Lates* can be recognized for management purposes such as mixed stock fisheries. Vij et al. (2014) note that the assumption of single species of Indo-Pacific *Lates* has undoubtedly influenced previous studies and microsatellite data from Thailand showing the highest allelic richness may reflect genotyping of two divergent genetic lineages along the west coast of Thailand (Yue et al., 2009). If different species of Indo-Pacific *Lates* are recognized, then questions of biology can be researched such as the causes of speciation and what the mechanisms for reproductive isolation of IND may be. Development of an anatomical key usable in the field that can separate these species e.g. (Wang et al., In Revision) would be of great practical utility and could be used to resolve the distributions and overlap of IND and SEA lineages. A high-throughput sequencing assay that can discriminate the main lineages could provide a rapid assessment of genetic lineage, hybridization levels and possible provenance in studies of food adulteration.

### Admixture

Overall, analyses that can quantify hybridization in this study are challenging because of the high innate diversity of the SEA lineage and limited sample sizes, notably that of the IND lineage (n = 13). The sequences were also obtained opportunistically from the NCBI Sequence Read Archive and not collected by the authors. With these cautions in mind, we find evidence of admixture between major genetic lineages within our data set as evidenced by the placement of Cambodia-origin samples, with a largely SEA genomic background within the mitochondrial IND lineage (**Figures 2, 3 & 4**). When examining all 60 individuals at *K* = 3, the two Cambodia individuals with IND mtDNA have ancestry coefficients (*Q*) of 0.29 and 0.43 with the IND genetic cluster, however, there is such a degree of variation in the SEA group with respect to admixture proportions and inconsistency across sampling locations that these measurements should not be accepted at face value. The Cambodia group has two individuals with Q = 0.00 for the IND genetic cluster and one with 0.17 for example when examining all individuals. Separate analysis of Cambodia and *L. calcarifer* IND individuals only show *Q* = 0.23 and 0.33 for the two Cambodia individuals with IND mtDNA and no evidence for nuclear genomic admixture of other Cambodian individuals with IND. This finding is congruent with the NewHybrids analysis indicating the presence of two F2 hybrids in the Cambodian fish, in line with the ancestry proportions of these fish analyzed with IND fish separately (**Supplemental Figure S7**).

Nucleotide diversity in the SEA Indonesia-SJ location is very high (**Figure 5B**) and no doubt influences the reduced F_st_ of this lineage to other sampling regions (**Figure 5A**). Combined with the large negative *D* value which arises from admixture (Hahn et al., 2013), this sampling location may contain admixed individuals. A similar pattern of high nucleotide diversity, low-F_st_, and strongly negative *D* is apparent with the Thailand sampling location (**Figure 5**). Both the Indonesia-SJ and Thailand sampling locations show substantial contributions of the IND genetic cluster in their ancestry (**Supplemental Figure S2**), but it is not uniformly distributed across sampling locations. If the distribution of ancestry is uniform across sampling locations, it would suggest an older age of hybridization. In NewHybrid analyses, Indonesia-SJ and Thailand contain numerous recent (F2) hybrids (Supplemental Figure S7). Furthermore, the admixture analyses suggest that Thailand at *K* = 4 is represented by more than one distinct genetic unit from within the SEA lineage with additional contributions from the IND lineage. The dearth of F1 hybrids found by NewHybrids was surprising but may have to do with a high degree of diversity within the SEA lineage as a result of (1) retained ancestral variation and (2) historical admixture. Furthermore, the selection of a random set of 200 SNPs due to computational limitations may not capture informative loci for hybridization assessment.

## Conclusions

We provide several advances in the study of Indo-Pacific *Lates*, presenting molecular phylogenetic analyses of a geographically and genomically representative data set and testing for adaptive differentiation and recent or more advanced hybridization. The analysis of genome-wide nuclear variants also provides evidence of hierarchical divergences, that is within the IND, SEA and AUS+NG lineages there is genetic structuring. The SEA lineage demonstrates a complex history with a combination of natural genetic differentiation and what is likely human mediated admixture events. Clearly, samples of known provenance, the use of informative loci that are diagnostic to lineages, and boosting of sampling locations and sample sizes focused across the possible intergrade zone IND and SEA would help resolve these ambiguities. Future studies making use of archive and museum samples to provide genetic baselines prior to use of BP as an aquaculture species can be used to create a genetic baseline to compare modern genetics against to identify potential translocations and adaptation to changing environments (Turbek et al., 2023).

Phenotypic and life history variation within Indo-Pacific *Lates* remain poorly characterized and are other aspects of variation that can substantially contribute to ecosystem productivity and commercial applications. Research indicates that within Australia for instance, there is strong local adaptation related to temperature (Newton et al., 2013) that could be applied to various goals. The wide and varied use of Indo-Pacific *Lates* in commercial, recreation and subsistence fisheries as well as aquaculture creates differing priorities across user groups. Further exploration of genotypic and phenotypic variation can support conservation aquaculture and food production as well as the preservation of productivity and adaptive potential of Indo-Pacific *Lates* stocks.

## Data Availability

Genomic sequence data are available from the NCBI Sequence Read Archive under BioProject accessions PRJNA311498, PRJNA1021005 and PRJDB13763. Alignments for phylogenetic analyses include the control block for SVDQuartets are included in the Data Supplement of this paper. Sequence data examined in this study for time-calibrated phylogenetic analysis are publicly available with mitochondrial sequence accessions reported in Table 1.

## Acknowledgements

The authors would like to recognize two anonymous reviewers that provided valuable feedback for improving the manuscript.

---

**Supplemental Figure S1.**
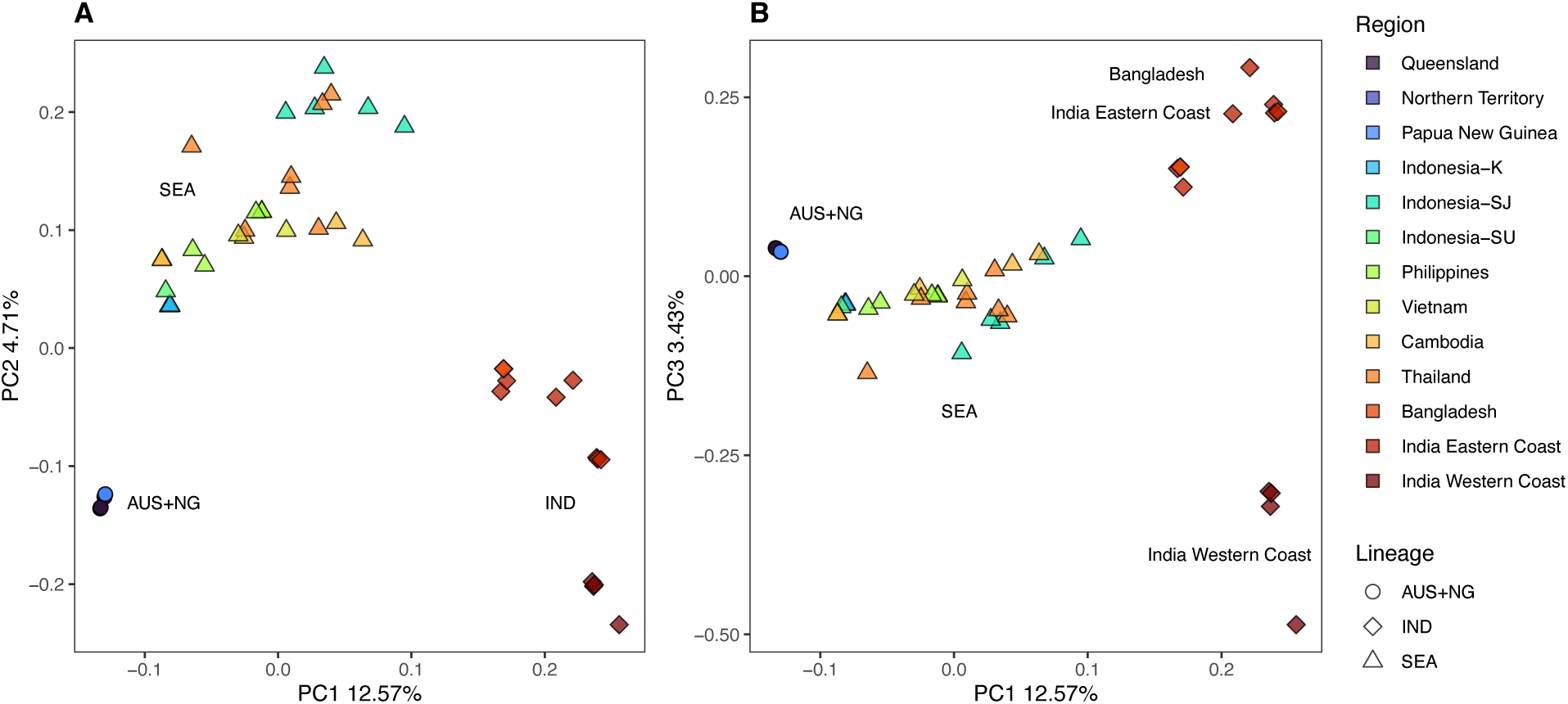
Principal Component (PC) analysis of the whole-genome sequence data set of 60 Indo-Pacific *Lates* utilizing 7,321,852 genotype likelihoods. The first PC is on the *x* – axis in both figures with the second PC on *y* – axis in (A) and third PC in (B).

**Supplemental Figure S2.**
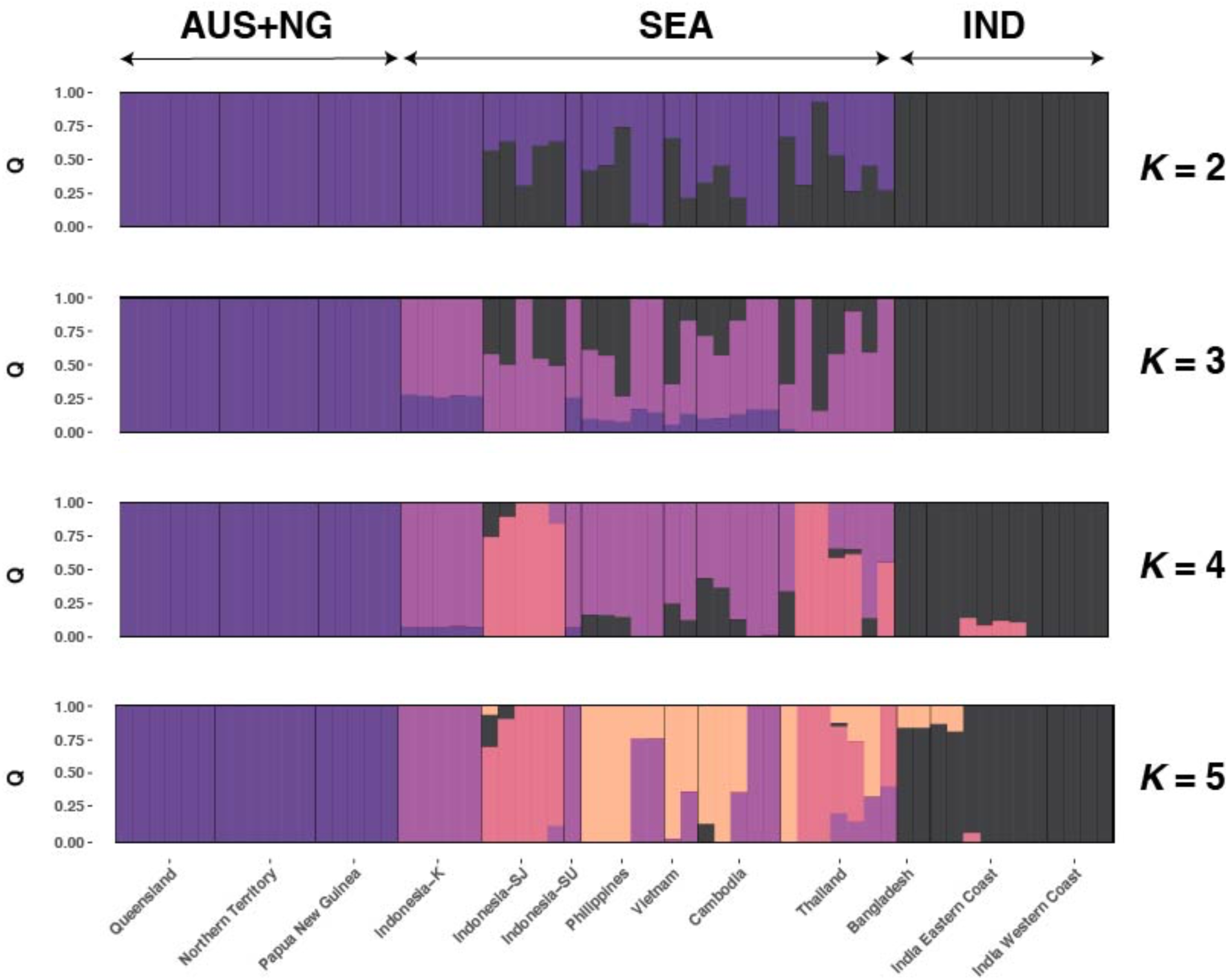
Genome-wide admixture analysis of Indo-Pacific *Lates* from WGS data (n = 60) from 7,321,852 genotype likelihoods for *K =* 2 - 5 genetic clusters. Individuals are plotted on the *x* – axis and ancestry coefficient (*Q*) on the *y* – axis.

**Supplemental Figure S3.**
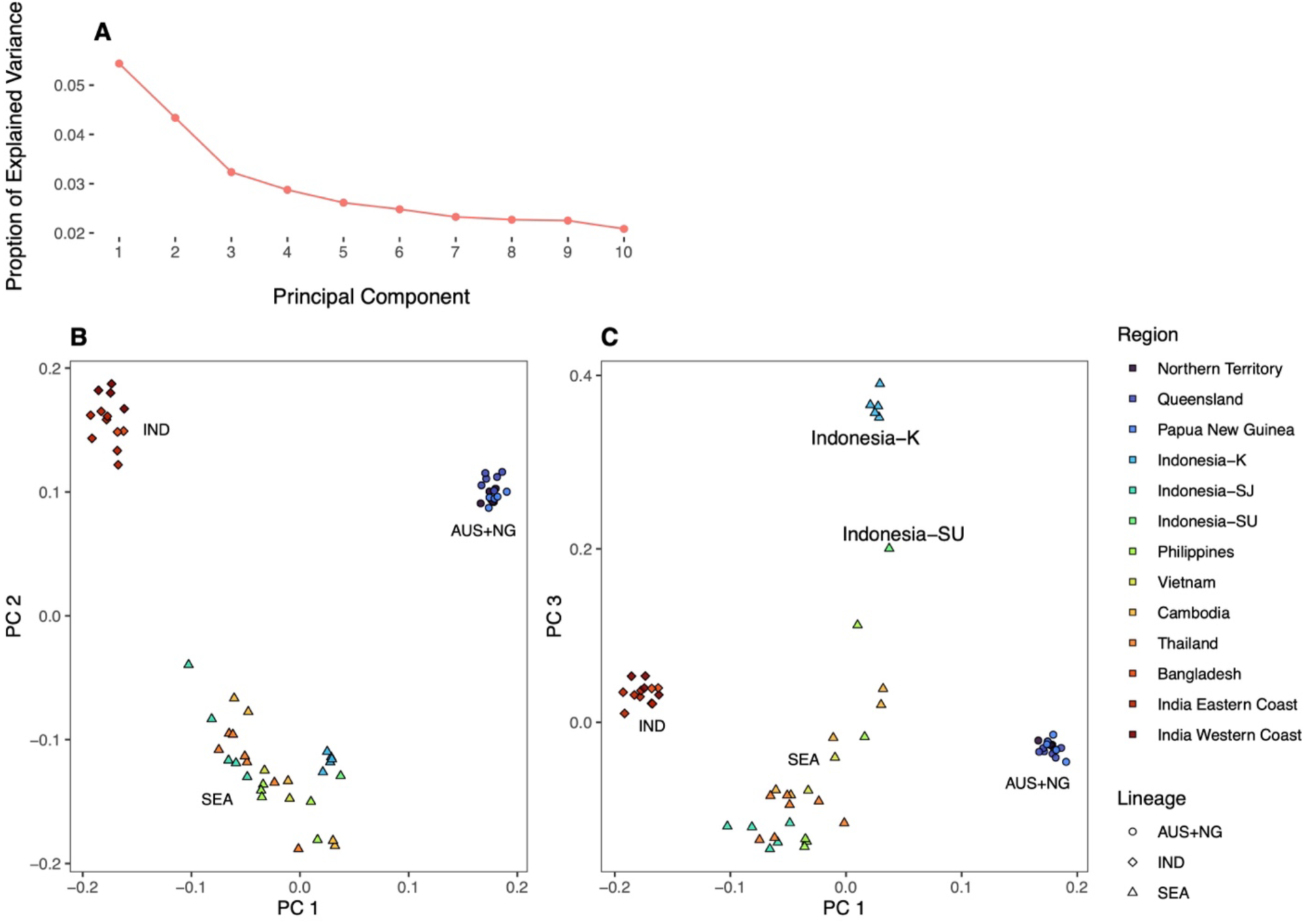
PCAdapt analyses for identification of adaptive genetic loci of all Indo-Pacific *Lates* analyzed in this study. **(A)** Proportion of explained variance for each Principal Component (PC), supporting *K* = 3 PCs. **(B)** Plot of PC 1 & PC 2, IND, SEA and AUS+NG. **(C)** Plot of PC 1 & PC 3, PC 3 separates Indonesia-K and Indonesia-SU sampling locations from all other samples.

**Supplemental Figure S4.**
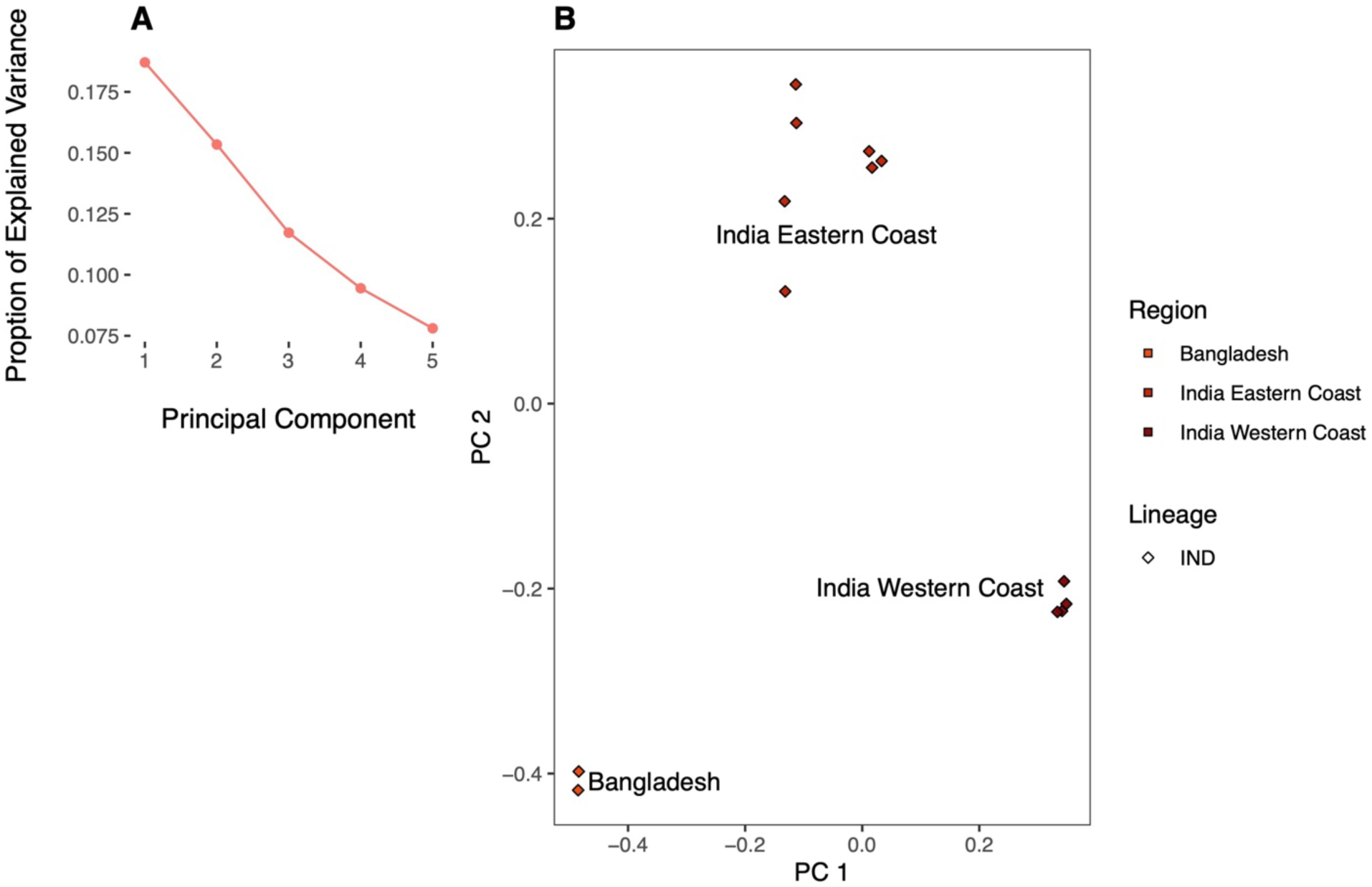
PCAdapt analyses for identification of adaptive genetic loci from the IND genetic lineage. **(A)** Proportion of explained variance for each Principal Component (PC). **(B)** Plot of PC 1 & PC 2.

**Supplemental Figure S5.**
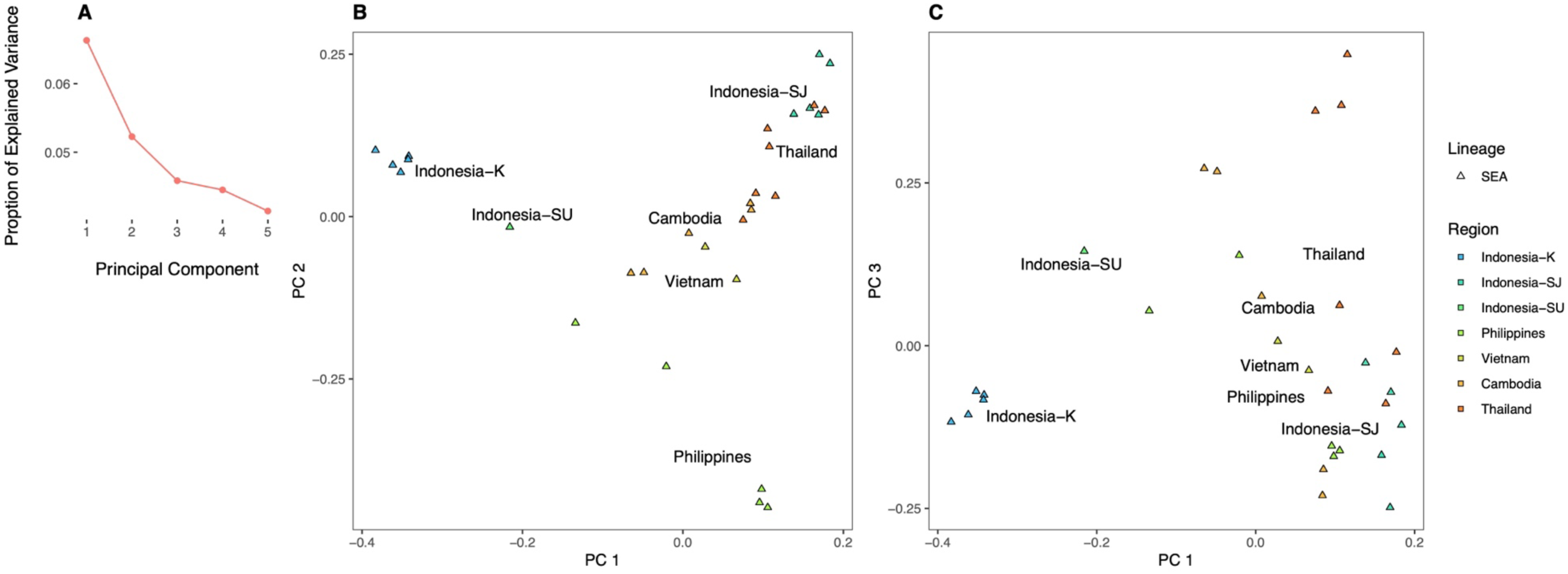
PCAdapt analyses for identification of adaptive genetic loci within the SEA lineage. **(A)** Proportion of explained variance for each Principal Component (PC), supporting at least *K* = 3 PCs. **(B)** Plot of PC 1 & PC 2, **(C)** Plot of PC 1 & PC 3.

**Supplemental Figure S6.**
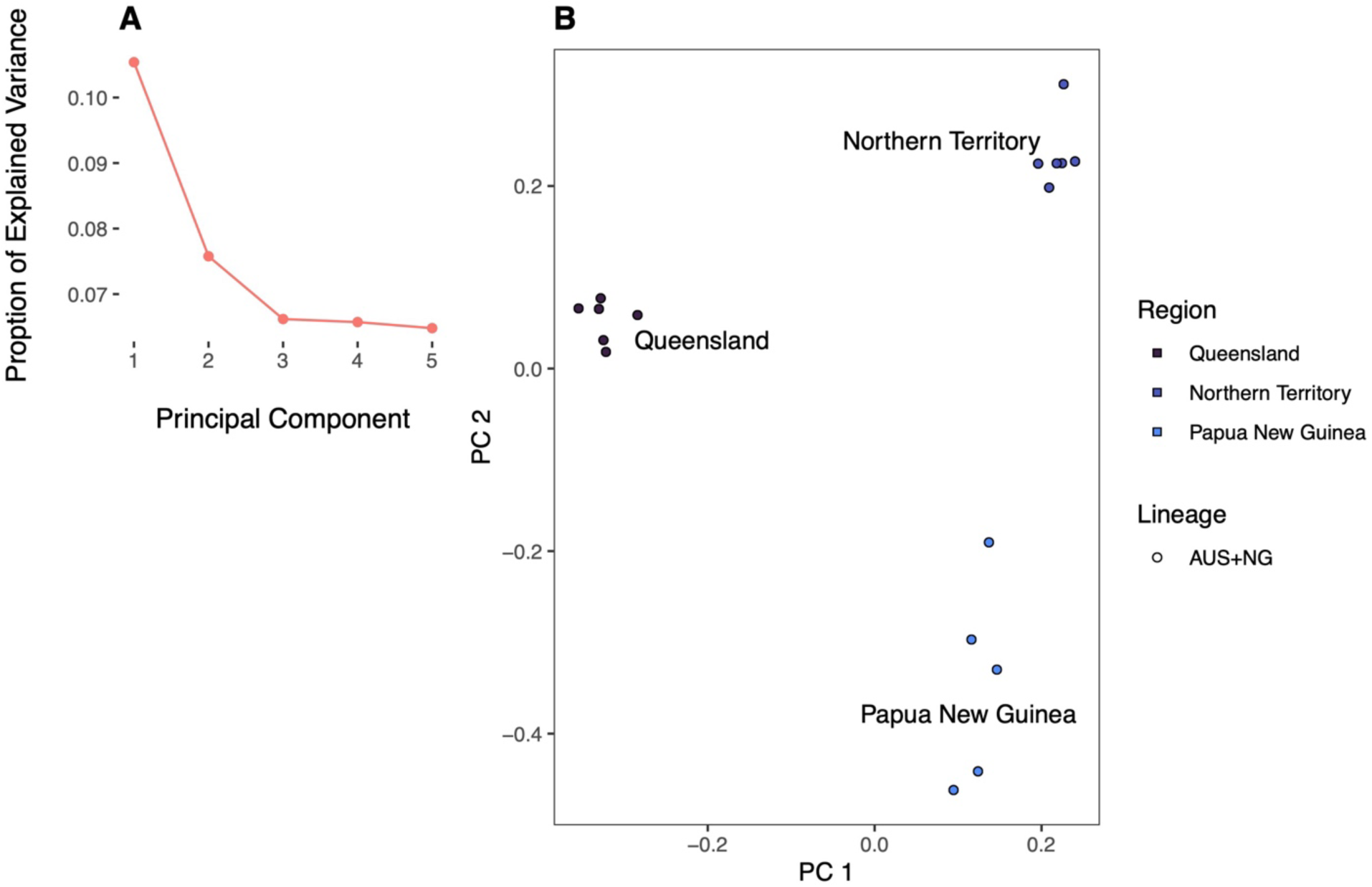
PCAdapt analyses for identification of adaptive genetic loci within the AUS+NG lineage. **(A)** Proportion of explained variance for each Principal Component (PC). **(B)** Plot of PC 1 & PC 2, separating Queensland, Northern Territory and Papua New Guinea sampling locations.

**Supplemental Figure S7.**
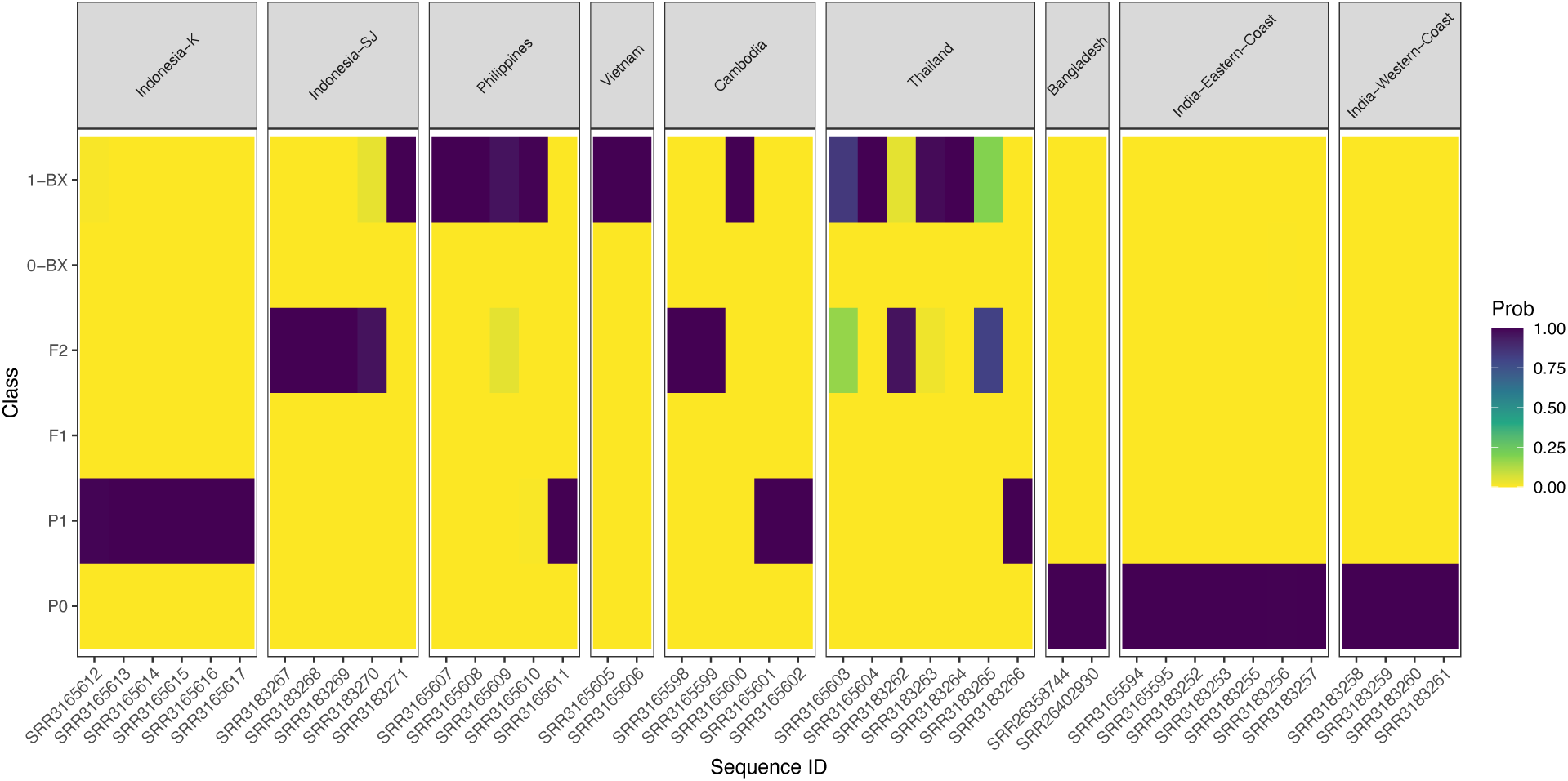
NewHybrids output from analysis of IND and SEA fish. For display, Indonesia-SU was combined with Indonesia-K. Individual sequence numbers, Sequence ID, are indicated on the *x* – axis. Parental populations, (P0, P1), recent hybrids (F1, F2) and backcrosses (0-BX, 1-BX) are indicated on the *y* – axis. Probability of assignment is indicated by a color gradient (Prob). Two individuals from Cambodia with IND lineage mtDNA are indicated to be recent hybrids (SRR3165998, SRR3165999).

